# Plants maximize competition while minimizing competitors belowground: a theoretical analysis of incentives for root competition in space

**DOI:** 10.1101/430504

**Authors:** Caroline E. Farrior

**Affiliations:** Department of Integrative Biology, University of Texas at Austin, Austin, Texas 78712

**Keywords:** Plant competition, adaptive dynamics, fine roots, resource limitation, tragedy of the commons

## Abstract

Recent research shows that shared access to belowground resources drives plants to overproliferate fine roots competitively, limiting community-level aboveground biomass. Models of this phenomenon are commonly based on an assumption that belowground resources and fine roots are thoroughly well mixed. In reality, of course, fine roots are spatially structured by individual. Here we investigate how costs of sending roots through horizontal space influence incentives for fine-root overproliferation. We find that these costs restrain overproliferation to the net benefit of community aboveground biomass. And further, the costs eliminate incentives for individuals to grow fine roots beyond their closest neighbors. Plants that interact with the fewest competitors benefit the most in relative fitness from overproliferation of fine roots. Effectively, individual-based optimization of root allocation in space increases the effects of competition while decreasing the number of individual competitors for each individual.

Because an individual’s optimal competitive network consists of only the closest neighbors, we predict the full effects of competition are achieved just shortly after disturbance, making competition belowground an almost inescapable pressure on plants. Together these results have important implications for predicting plant interaction networks, patterns of carbon allocation, and ecosystem carbon storage.

## Introduction

Understanding the relative importance of the environment versus neighboring individuals on an individual’s success has long been of great interest to ecologists (Connell 1983; Schoener 1983; Fowler 1986; Wilson and Tilman 1991; Goldberg and Barton 1992; Peltzer et al. 1998; Goldberg et al. 1999). Few concepts in ecology are as classic and appealing as the concept of a “ghost of competition past” (Connell 1980). Individuals with strategies that free themselves from competition with other species should be favored, and, over time, lead to filling of niche space by a diversity of species. Diversification by niche differentiation is not always possible, however. Limitation of essential resources like light, water, and nitrogen are challenges common to all plants. And, even if niche diversification is successful, it cannot free individuals from competition with other individuals of the same species. When individuals share access to the same pools of limiting resources, selection at the level of the individual often leads to intensification of competition through “red-queen” and “tragedy of the commons” dynamics (Hardin et al. 1968; Gersani et al. 2001).

One of the most well-recognized products of this intensification of competition is overproliferation of fine roots by individuals competing for water and nutrients with one another in horizontal space (Gersani et al. 2001; Maina et al. 2002; Zea-Cabrera et al. 2006*b*; Semchenko et al. 2010). But, our understanding of its determinants and variation is still in its infancy. Models of belowground competition often rely on an assumption that all individuals share access to the same resource pool belowground, one that is well-mixed across space or “mean-field” (Gersani et al. 2001; Zea-Cabrera et al. 2006*b*,*a*; Kiniry et al. 2008; Farrior et al. 2013*a*,*b*, but see Tilman 1994; Zea-Cabrera et al. 2006*b*; O’Brien et al. 2007). In reality of course, at some local scale, a root from one individual will interact more closely with its own roots than with the roots of other individuals. The roots must converge in space before connecting to a central stem which then runs aboveground. Access to belowground resources cannot be completely well mixed. And the importance of this fact may vary across gradients in productivity and individual density.

Although many experiments show plants respond to experimental manipulations of their competitive context by increasing fine roots and decreasing allocation to reproduction, many species do not show this or any response (Smyčka and Herben 2017). Some have raised the idea that this might mean those species do not engage in competition belowground as the others do (McNickle and Brown 2014). So, do individuals always have incentives to seek competition with one another, belowground? Do the incentives for belowground competition change across resource gradients as individual density declines? Competitive arms races, have the potential to drive big changes in vegetation structure and may play a role in the distribution of forests, grasslands, and shrublands (Staver et al. 2011; McNickle et al. 2016).

Here we develop two simple, but individual-based models with a consideration of discrete individuals and their space belowground to address these questions. I use individual-based optimization, or adaptive dynamics, to determine the competitive-dominant allocation to and placement of fine roots in horizontal space. With these models, we find that, only in the extreme case when the cost of a fine root does not depend on its distance from the stem, do we recover the mean-field model. If the cost of a fine root increases with its distance from the stem, we find competitive individuals have a preference for home soil, competitive overinvestments are restrained, and an increase in aboveground productivity results. We find that the competitive overinvestments are strongest when there are only two individuals competing. If many individuals are involved in competition, the effects of one individual on the others diffuses and weakens. As such, when individuals are placed on a spatial grid, we find the best strategy for an individual is to keep its roots close to home, competing with the fewest individuals possible and thereby increasing overinvestments in competition – maximizing the effects of competition while minimizing competitors. This strategy has the highest payoff to the individual and dominates across a wide range of environments.

## Materials and Methods

Here I describe two models designed to investigate the nature of competition among individuals for water and nutrients through horizontal space. The first model serves specifically to disentangle the effects of the number of competitors from resource availability on incentives for competition only. The second model builds on the first, placing the individuals in a more realistic spatial context and incorporating population dynamic feedbacks.

Both models are as simplified as possible to allow us to clearly understand model results and to develop new hypotheses about belowground competition without missing critical mechanisms. All plants are assumed to be exactly the same except for their allocation to and placement of fine roots so that we may understand how these strategies interact with the environment in the absence of variation in other phenomena.

Table 1 lists the variables and parameters used throughout the paper.

**Table 1:** Model symbols and definitions, in order of appearance

| Symbol | Description | Units | Default value |
| --- | --- | --- | --- |
| $n$ | number of individuals competing | ind | 3 |
| $a$ | area of one individual's home space | $\text{dm}^{-2}$ | 1 |
| $k$, subscript | individual identity | NA | |
| $A$ | Photosynthetic rate | $\text{gC dm}^{-2} \text{yr}^{-1}$ | |
| $l$ | leaf biomass | $\text{gC}$ | |
| $r$ | fine-root biomass | $\text{gC}$ | |
| $f$ | fecundity | $\text{ind yr}^{-1}$ | |
| $x$ | proportion of fine roots sent "away" | [unitless] | |
| $c_l$ | leaf costs* | $\text{gC gC}^{-1} \text{yr}^{-1}$ | 3 |
| $c_r$ | fine root costs* | $\text{gC gC}^{-1} \text{yr}^{-1}$ | 1.7 |
| $c_f$ | cost of reproduction* | $\text{gC ind}^{-1}$ | 1 |
| $c_x$ | cost of sending fine roots "away" | $\text{gC gC}^{-1} \text{yr}^{-1}$ | 1.5 (Model 1) |
| $w_k$ | water uptake by individual $k$ | $\text{L yr}^{-1}$ | |
| $\omega$ | intrinsic water-use efficiency | $\text{gC L}^{-1}$ | 2 |
| $u_S$ | water uptake efficiency of fine-root biomass | $\text{gC}^{-1} \text{yr}^{-1}$ | 1 |
| $S_k$ | water available at the home site of individual $k$ | L | |
| $R$ | rainfall | $\text{dm yr}^{-1}$ | 2 (or 20 if $q = 1$) |
| $\Delta$ | first-order rate of water lost to leakage | $\text{yr}^{-1}$ | 0.02 |
| $r_{T,k}$ | total fine-root biomass at home site of individual, $k$ g | $\text{gC}$ | |
| $\mu$ | mortality rate | $\text{yr}^{-1}$ | 1 |
| $n_d$ | site-level individual density | $\text{ind dm}^{-2}$ | 1 (Model 1) |
| $c_{x,0}$ | soil specific cost of placement of fine roots | $\text{gC gC}^{-1} \text{dm}^{-2} \text{yr}^{-1}$ | 1.2 (Eq 8, Model 2) |
| $q$ | proportion of growing season without water limitation | [unitless] | 0 |
| $N$ | net-nitrogen mineralization rate | $\text{gN dm}^{-2} \text{yr}^{-1}$ | 0.003 (or 0.204 if $q = 0$) |
| $V$ | maximum photosynthetic rate per leaf area | $\text{gC dm}^{-2} \text{yr}^{-1}$ | 8 |
| $\sigma$ | leaf mass per unit area | $\text{gC dm}^{-2} 0.3$ | |
| $\alpha_f$ | leaf-level photosynthetic dependence on light | $\text{gC MJ PAR}^{-1}$ | 1.81 |
| $L_0$ | photosynthetically active radiation | $\text{MJ PAR dm}^{-2} \text{yr}^{-1}$ | 12 |
| $k$ | light extinction through the canopy | [unitless] | 0.5 |
| $\rho$ | whole-plant nitrogen demand per unit leaf biomass | $\text{gN gC}^{-1} \text{yr}^{-1}$ | 0.107 |

Note: dm stands for decimeters. *Costs include annualized building, maintenance and respiration costs. Cost of reproduction includes the cost of all reproductive structures and seeds needed to produce one new individual. See Farrior et al. (2013*b*) for explanations and derivations of parameter values.

### Model 1, “n - competitor model”

In this first model, the number of individuals with the potential to share resources is taken to be a constant, *n* (ind). Assume each individual (distinguished by subscript, *k*) takes up the same amount of space (*a*, 10 cm x 10 cm or 1 dm^2^) and each unit of space has the same environmental inputs (light, water, and nitrogen). That is, given specific plant traits, changing *n* does not change an individuals’ resource availability.

For simplicity, also assume the plants are annuals. Plants fix carbon through photosynthesis (*A_k_*, gC yr^−1^ and that carbon is used to pay for the building and maintenance costs of plant tissues: leaves (*l_k_*, gC), fine roots (*r_k_*, gC), and reproduction ( *f_k_*, ind yr^−1^). 
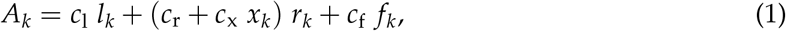

Where *c*_1_ (gC gC^−1^ yr^−^1), *c*_r_ (gC gC^−1^ yr^−1^), and *c*_f_ (gC ind^−1^) are the annualized costs of building and maintenance of leaves, fine roots, and new individuals (fecundity), respectively. An additional cost, *c*_x_ is applied to the proportion of roots that are sent a greater distance to compete with other individuals (*x_k_ r_k_*).

For each individual in this model, there are two types of space belowground: “home” space and “away” space (figure 1). We define home space as all of the space that is closer to an individual than to any of the other individuals. Away space comprises the home spaces of all of the other (*n* − 1) individuals. All space in the model is the home space of one and only one individual. Despite the physical impossibility, we assume that the home spaces of competitors are all equally close to one another. In model 2, we relax this assumption.

**Figure 1:**
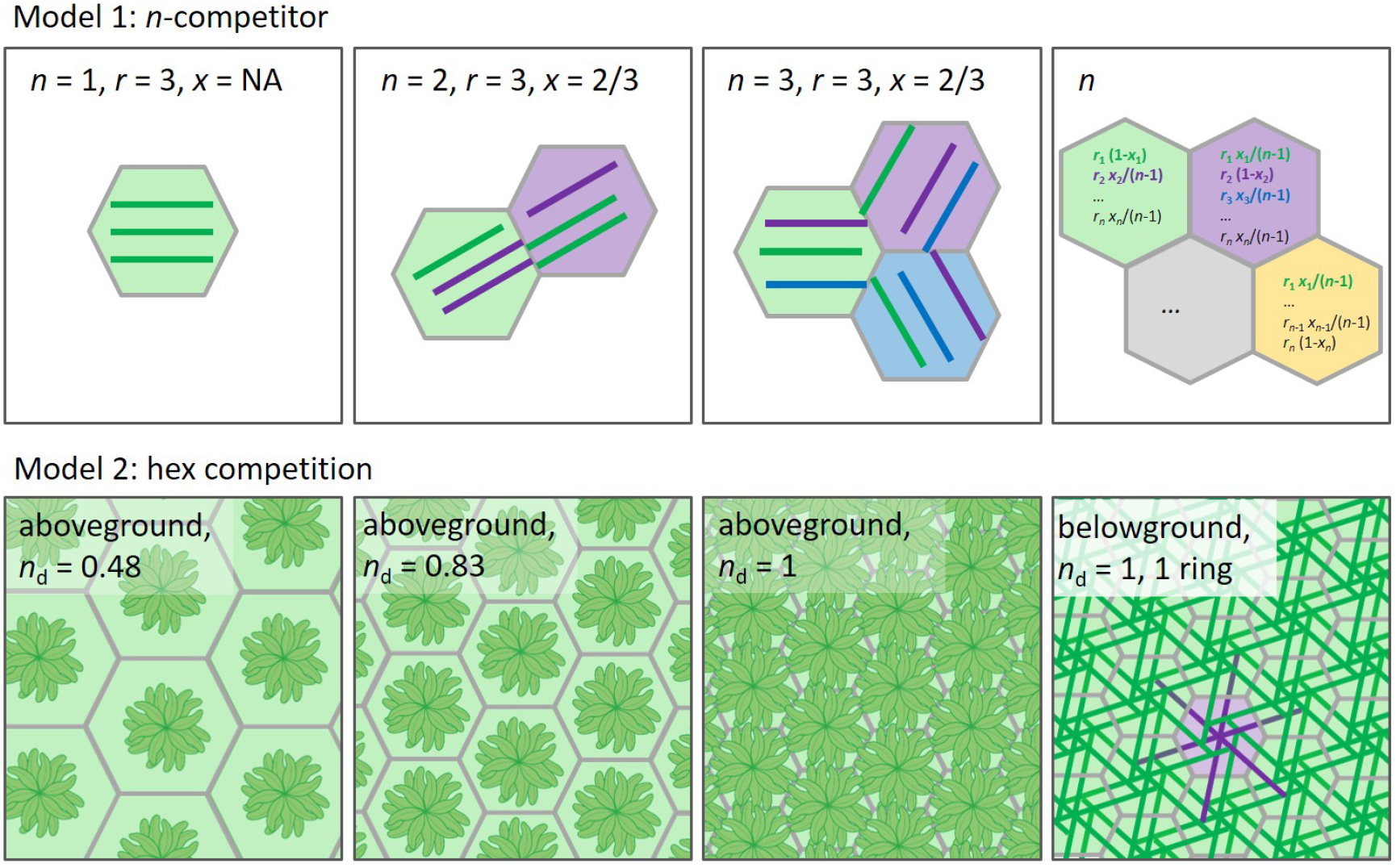
Illustrations of Model 1 (top panels) and Model 2 (bottom panels). Each hexagon represents one individual’s “home space” and each individual’s roots are represented by lines matching the color of their home space. **Model 1:** Each individual has the same amount of space and resources and is equally connected to every other individual. Across the first three panels, we see the effect of changing *n* if all individuals have the same strategy of fine root investment (*r*) and placement (*x*, the proportion sent to compete with neighbors). The final panel shows the general case of *n* individuals, each with their own strategy (*r_k_* and *x_k_*). Model 2: A portion of the 96 individual hexagonal torus is depicted. Individual density (*n*_d_) increases across the first three panels. As belowground home space decreases with density so too does the cost of sending roots toward neighbor a set number of neighbors (6, 12, or 18; *c*_x_(# rings), eq. 8). The final panel depicts fine roots belowground for a case in which a target individual (purple) is competing with its 6 closest neighbors (green). As belowground competition is the focus of this theoretical investigation, in all cases leaf area per individual is constant.

The rate of photosynthesis for an individual (*A_k_*) may be limited by light, water, and/or nitrogen. For efficiency, in the main text, we describe only the case of constant water limitation. Appendix A provides physiological details for the cases of light and nitrogen limitation. All of the formulations follow Farrior et al. (2013*b*).

For water-limited plants, photosynthetic rate it proportional to water uptake (*w*, L yr^−1^): 
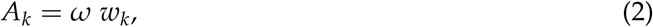
where *ω* is the water use efficiency (gC L^−1^). The uptake of water by individual *k* (*w_k_*) depends on the available water at both home (*S_k_*, L) and away sites (*S_j_*, where *j ≠ k*):
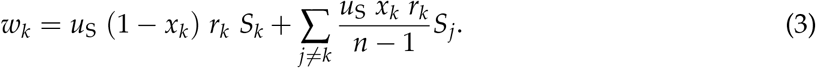

Where (1 *− x_k_*) is the proportion of the individual’s *r_k_* fine roots that stay at home and *u*_S_ is the uptake efficiency of fine-root biomass for water (gC^−1^ yr^−1^).

Water available at a site is the balance of incoming rainfall, *R* (dm yr^−1^), first-order soil-moisture dependent losses from the rooting zone (Δ, yr^−1^), and uptake by all plants with fine roots at the site (*r*_T,*k*_, eq. A1): 
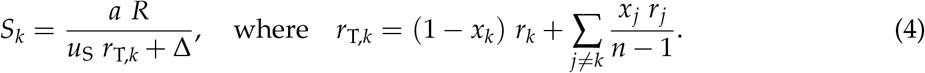

Note, we assume horizontal movement of water across sites is negligible. Although water moves well through soils, at some scale there is spatial structure to water uptake by individuals. To study this in the simplest case, we assume here that water movement from one individuals’ home space to the next is not significant. The parameter *a* is the area of one home site (here 1, in model 2: 1/*n*_d_, dm^2^).

In the case of constant water limitation by definition, all leaf biomass needed to achieve the water-limited photosynthetic rate is more valuable to the plant than it costs. Competitive plants then will hold exactly this amount of leaf biomass (*l_k_*), implicitly defined: *A*_L_(*l_k_*) = *ω w_k_*, where *A*_L_ is the photosynthetic rate for a plant canopy given light level and leaf biomass (eq A4).

Given, *r_k_, x_k_*, and *l_k_*, we can solve the individual carbon conservation equation (eq. 1) to find the individual’s allocation to reproduction: 
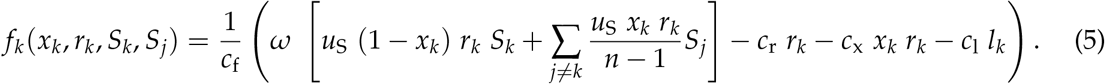

Because we assume mortality rate to be independent of the plant strategies of investigation, this allocation to reproduction is treated as a measure of individual fitness.

### Model 2, “hex-grid competition model”

Here we will expand on Model 1 by adding an explicit treatment of space and include a feedback between allocation to reproduction and the density of individuals (*n*_d_, ind dm^−2^). We maintain the assumption that, given population density, all individuals occupy equal amounts of space (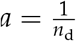, figure 1). We use a population size of 96 individuals and arrange individuals on a hexagonal grid whose spatial scale changes with density. The grid is wrapped on a torus to avoid edge effects.

Individual density in this model is the result of the reproduction ( *f_k_*) and the mortality (*μ*, yr^−1^) of individuals in the site: 
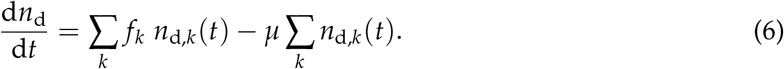

The resulting density can be found by solving the above at equilibrium. If all individuals in the site have the same strategies reproduction will simply balance mortality: 
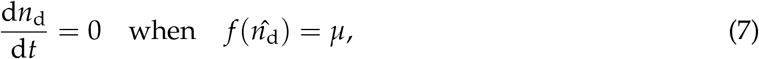
where 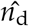 is the equilibrium individual density of the site.

As density changes, the distance between individuals changes and thus the cost of sending roots to another individuals’ home space should change. We assume the cost of roots sent away increases with the square of the distance over which plants send their roots: 
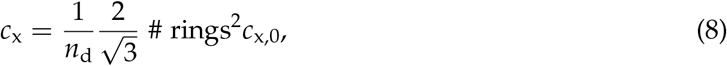
where *c*_x,0_ is a constant property of the soil and plants (gC gC^−1^ dm^−1^ yr^−1^), “# rings” is the number of concentric rings of hexagons across which an individual spreads its *x_k_* * *r_k_* roots. Individuals can send their roots different distances but must compete with all individuals in a ring if they compete with any in a ring. For example, if roots extend to the first ring, the individual competes with 6 individuals, to the second ring 18 individuals, etc… We simply assume all roots sent away are distributed evenly among the away spaces, independent of their distance to the individual. Other functional forms for the distribution of roots across distance and the dependence of cost on that distance were considered. These changes had no qualitative effect on results, however.

Because we are focused here on understanding belowground competition, we assume the light-intercepting area (“crown area”) of the plants is a constant (1 dm^2^, represented by cartoon leaf area in figure 1, bottom panels). Further we analyze the model only for conditions under which the density of individuals is low enough that all aboveground biomass is in full sun (*n*_d_ = 1, figure 1).

### Analysis

With the frameworks for both the “*n*-competitor” (Model 1) and the “hex-grid competition” (Model 2) models, we search for belowground plant strategies that will dominate at equilibrium. These strategies may be achieved through individual plasticity, community assembly, or adaptive trait evolution. We consider fine roots (*r*), the proportion of fine roots plants send away (*x*), and the number of rings of competitors they send their roots to (Model 2 only) as traits. Leaf biomass *l* is also taken to be value that optimizes fitness.

We follow an evolutionary game theoretic or adaptive dynamics approach (Maynard Smith and Price 1973; Maynard Smith 1982). We search for strategy sets that, when dominant at the site (resident), cannot be invaded by other strategies and are achievable through mutations of small effect (ESS and CSS, Geritz et al. 1998).

Many analyses like this are based on models with an infinite number of individuals and it can be assumed that the mutant has a negligible effect on the residents’ environment. That is not the case here. We assume one individual has the mutant strategy and the rest of the individuals have the resident strategy. For example, in the first model, this gives us the following fitness functions for the resident (R) and mutant (M) strategy types, respectively: 
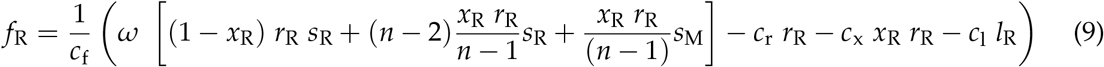
and 
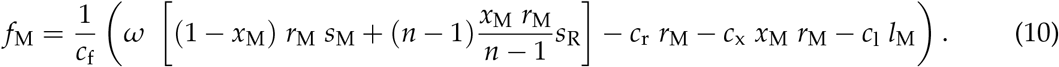

The competitive dominant, the evolutionarily stable strategy, if one exists, is the strategy that cannot be invaded by any other. It satisfies the following (following Maynard Smith and Price 1973; Maynard Smith 1982): 
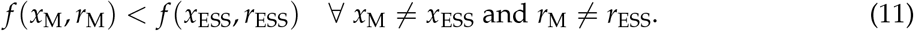

Because the influence of the strategy of the mutant on the environment is significant here, solutions of the ESS sets must be found numerically. A generic method for finding putative two-dimensional ESS sets numerically has been written for this project. This code and all other code necessary for this paper can be found online (code written in R, R Development Core Team 2008, https://github.com/cfarrior/comp-terr-VP). Plots verifying solution sets are both evolutionarily and convergence stable are provided for a representative sampling of the results (Appendix B).

To find the ESS sets including the third trait, the distance to send roots (Model 2 only), a few more details are needed. We compete discrete strategies against one another: 0 competitors, 6 competitors (1 ring), and 18 competitors (2 rings). Where the number of competitors is a trait of the individual: this is the number of individuals that an individual send its roots to, not necessarily equal to how many individuals are sending roots back to its home space. Because the specific values of *r* and *x* (resident ESS or invader ESS) did not change the results of this pairwise competition, values of 1 and 0.5, respectively were used.

## Results

We begin with results from Model 1, the “*n*-competitor model” designed to investigate the role of the number of competitors on incentives for competition belowground.

### Cost of competition increases community fitness

If there are no costs of sending roots to neighbors’ home soil (*c*_x_ = 0, no cost of sending roots through space), a classic mean field model of belowground competition matches the results of our competitive optimization analysis. Plants equally engage in the full overinvestment in fine-root biomass for competition if they have an infinite number of competitors or if there is only one competitor.

The following exact predictions for a mean field case match numerical results for the case of *c*_x_ = 0 (figure 2, gray lines versus dots): 
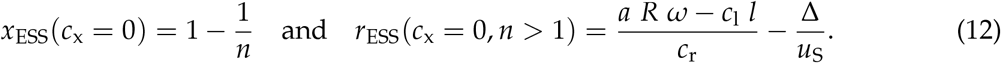

**Figure 2:**
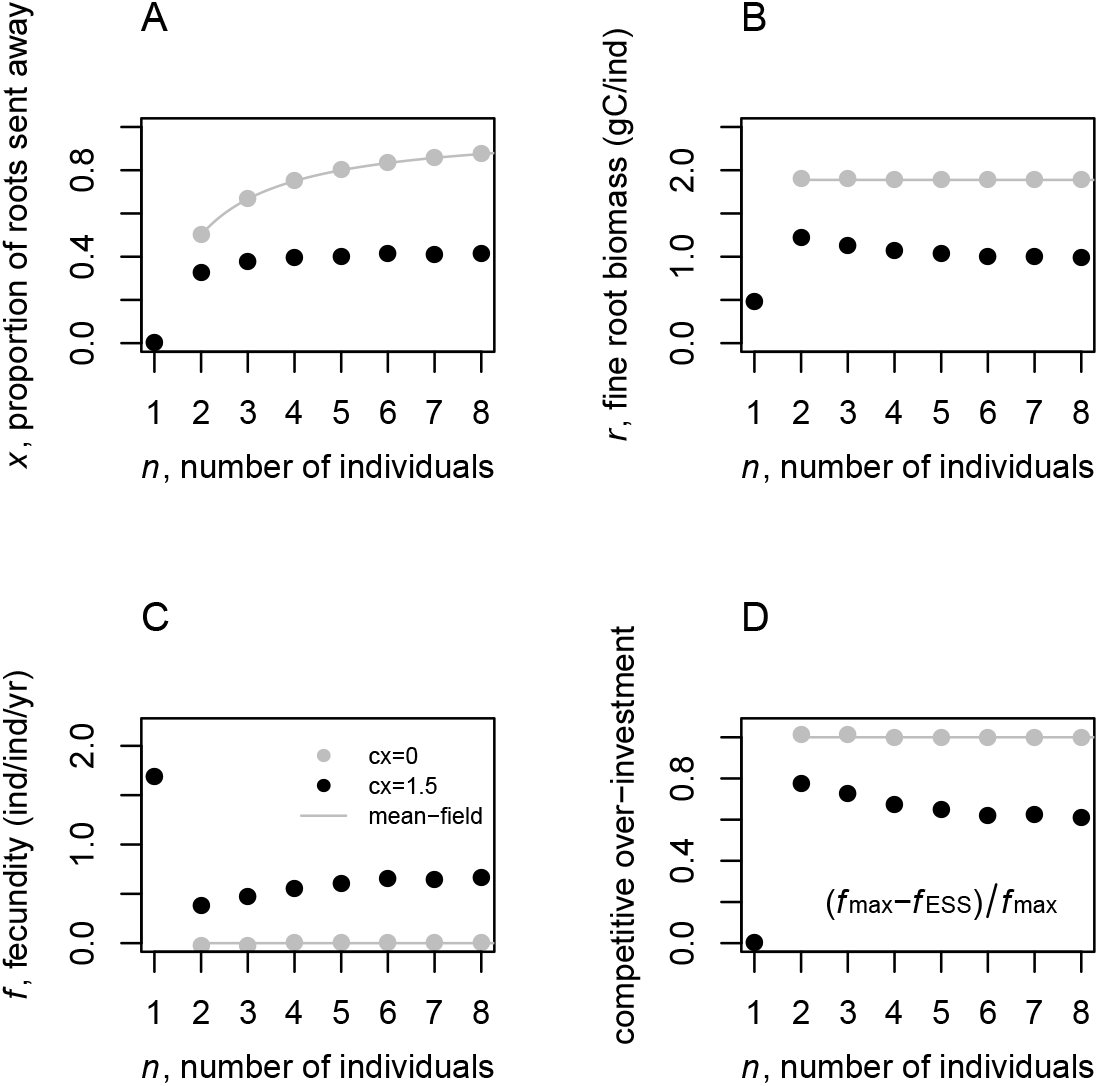
The effect of the number of individuals sharing belowground resources (*n*) on competitive dominant (ESS) fine-root allocation and placement (Model 1). (A) The proportion of fine roots sent to neighbors’ home space (*x*_ESS_), (B) fine-root biomass (*r*_ESS_, gC ind^−1^), and (C) reproductive output ( *f*_ESS_, ind ind^−1^ yr^−1^) of the competitive-dominant strategy (ESS) set. (D) The fraction of potential reproductive output diverted to competitive overinvestments, here fine roots. Results shown are for cases of zero (gray) and non-zero (black) costs of extending roots to neighbors’ home space (*c*_x_). Lines show the predictions for an even distribution of fine roots (A) and well-mixed or mean-field water availability (B-D, eq. 12). If not otherwise indicated, parameter values are those found in Table 1.

This strategy results in a “full” tragedy of the commons, the ESS strategies in monoculture have no allocation to reproduction as in Zea-Cabrera et al. (2006*b*).

If the cost of competition, *c*_x_, however is positive we find qualitatively different results. Investment in fine roots (*r*) and in sending fine roots away (*x*) decreases with *c*_x_. This occurs as well when we increase the cost applied to all fine-root biomass (*c*_r_). But, with a change in *c*_r_ alone, there is no net change on the plants’ allocation to other tissues. When *c*_x_ increases, there is a net benefit on community-level productivity in allocation to reproduction.

### Incentives for fine-root overproliferation are highest when there are the fewest competitors

As in other studies, we find that by adding the possibility of more than one individual sharing access to resources, there is an increase in allocation to fine roots (*r*) to the net detriment of allocation to reproduction at equilibrium ( *f* (*n* = 1) to *f* (*n* = 2), figure 2C). But, by adding additional individuals (if *c*_x_ > 0 and *n* > 2), we find that this overproliferation of fine roots decreases and allocation to reproduction increases (figure 2B,D).

Now we move to Model 2 to find the implications of these results within a spatially explicit context while incorporating population dynamics.

### Competitive dominant plants intensify competition by minimizing competitors

From an evolutionary analysis of *r, x*, and the number of competitors to interact with in space, we find that the competitive-dominant strategy is almost always to compete with as few individuals as possible, while remaining in competition (here 6, figure 3). From our results of Model 1, we know this reduction in effective competitors has the effect of increasing the payoffs to overproliferation of fine roots (figure 2B, highest root biomass with two competitors).

**Figure 3:**
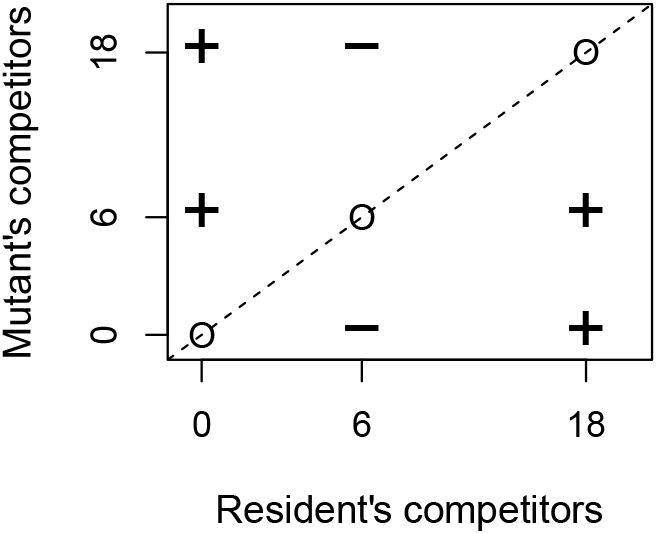
The results of pairwise competition among individuals with differing strategies of fine root placement (Model 2, hex-competitor model). Individuals can keep roots at home (0 competitors), send them to their closest ring of neighbors (6 competitors), or send them to the closest two rings of neighbors (18 competitors). All individuals in the 96 individual grid have the resident strategy except for one mutant. (+) marks cases in which the mutant strategy performs better than the average resident 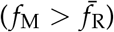, (−) marks cases in which the average resident performs better than the mutant 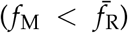, and (0) indicates neutrality 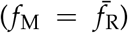. Here, we see the strategy of competing with only closest neighbors (6) is the competitive dominant (ESS). It cannot be invaded by any other strategy when resident. This result was found for a broad range of environmental conditions (including all cases where *c*_x_ > 0 and *x*_ESS_ ≠ 0, see main text for further detail).

Only when there are no costs of competition (*c*_x,0_ = 0) is competing with more than the fewest possible individuals a good strategy. The strategy of no competition, keeping all roots at home (*x*_ESS_ = 0) is only the best strategy when plants are investing so many fine roots to forage in leaky soil that competition with one another is unimportant. That is, Δ is high, *c*_x,0_ is high, and either *R* or *N* is low if *q* is low or high, respectively (figure 4). If leakage (Δ) is zero, however, there are no conditions under which the most competitive strategy is to keep roots at home (*x*_ESS_ is never zero, figure C3). When the competitive strategy is for roots stay at home (*x*_ESS_ = 0) the community-level optimal allocation to fine roots is equal, of course, to the ESS allocation (figure C5).

**Figure 4:**
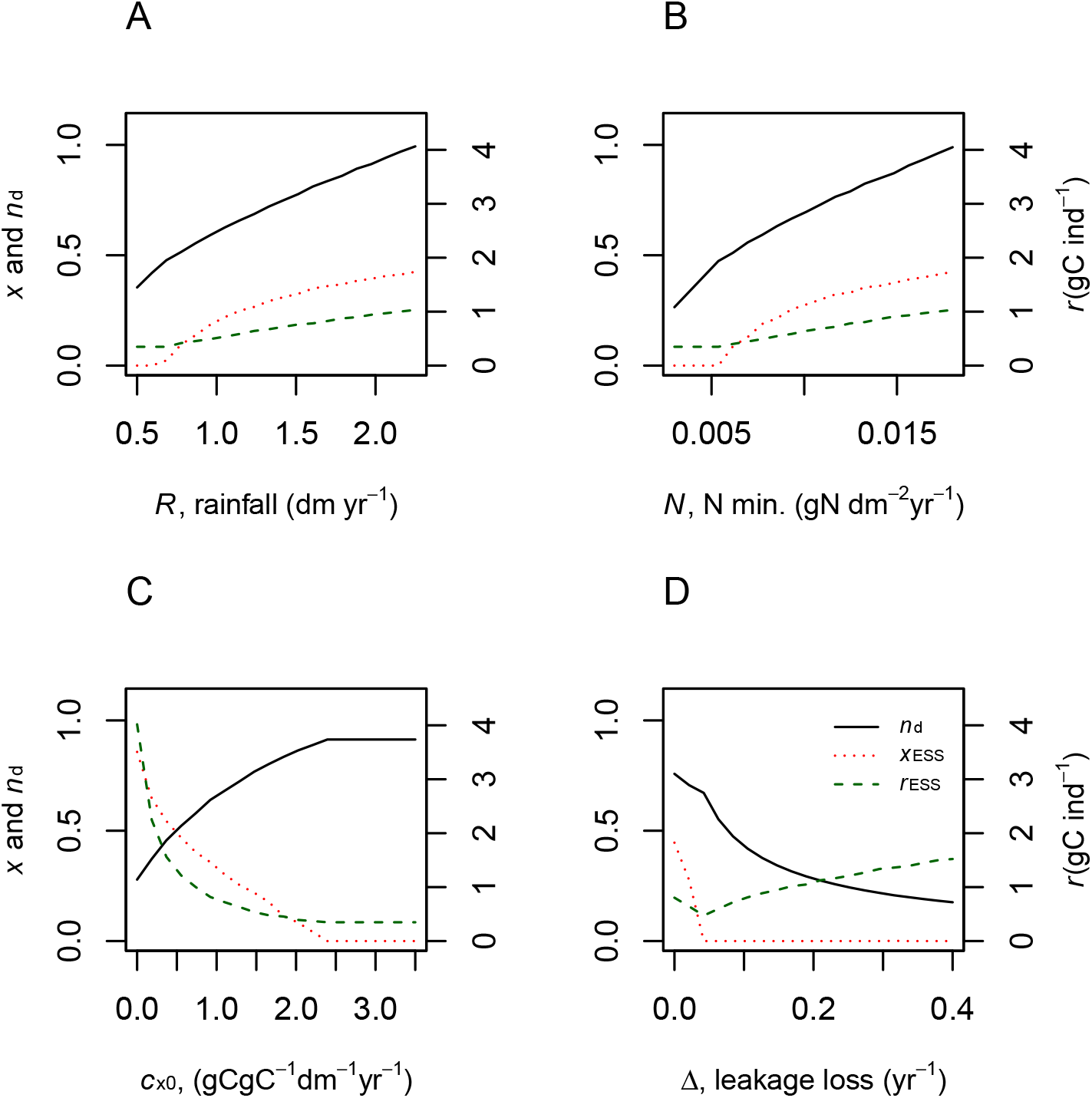
The effects of belowground competition across environmental gradients: (A) *R* rainfall, (B) *N* nitrogen-mineralization rate, (C) *c*_x,0_, the cost of extending roots through space (eq 8), and (D) Δ the first-order leakage loss rates of water and nitrogen from the soil. Each panel shows results for fine-root biomass (*r*_ESS_, gC ind^−1^) on the left axis and proportion of fine roots sent to compete with neighbors (*x*_ESS_) and resulting individual density (ind dm^−2^) on the right axis. The proportion of time spent with enough water to not be water limited is *q*, a parameter set to 0, 1, 0.5, and 0.5 in A through D, respectively. Values of all other parameters can be found in Table 1.

### Strength of competition across environmental gradients

Plants respond to increasing rainfall or nitrogen by increasing individual density and increasing individual allocation to fine roots (4). At the lowest resource levels (low *R* or *N*), we find that plants keep all of their roots at home. Again, this is not the case if resource dependent leakage losses (Δ) are zero (C3). Whenever *x* is greater than zero, the best strategy for these plants is to compete with only the closest neighbors (figure 3). The number of competitors they engage with does not change when plants are in competition, but we do see the relative importance of competition change as the density of individuals increases. As resource inputs increase and density follows, the cost of competing with the closest neighbors decreases (eq. 8) and the proportional allocation belowground to competition (*x*_ESS_) increases.

Both increasing the extrinsic cost of root extension through space (*c*_x,0_) and rate of first-order leakage losses of belowground resources (Δ) decrease the allocation to fine roots. But, their effects on aboveground biomass differ. Increasing *c*_x,0_ frees plants from competition as increasing *c*_x_ in Model 1 did and results in an increase in individual density. In contrast, increasing the rate of leakage losses (Δ) is accompanied by a decrease in aboveground productivity.

## Discussion

I have presented here two models distilling the incentives for and effects of competition among individuals across space. We have found a number of insights into and predictions for general plant strategies of competition.

Only if there are no costs to competition – specifically, no costs of extending roots closer to competitors than oneself – do we find that plant communities satisfy the assumptions of a mean field approximation. In this extreme case, individuals invest in so many fine roots for individual gain that at ecological and evolutionary equilibrium the community has no carbon left to invest in reproduction, the “paradox” of the tragedy of the commons (Zea-Cabrera et al. 2006*b*). We have found before that this paradox is resolved if one considers a realistic amount of variability in water availability (Farrior et al. (2013*a*)). But, we find a completely different resolution here by including the cost of extending fine roots through horizontal space.

With costs of sending some roots to forage closer to neighbors than oneself (*c*_x_ nonzero), the drive to send roots towards competitors is diminished (*x*_ESS_ decreases) as are the incentives for fine-root overproliferation (*r*_ESS_ decreases). If *c*_x_ increases, the decrease in allocation to fine-root biomass (*r*_ESS_) more than compensates for the new cost, resulting in higher allocation to reproduction at evolutionary equilibrium. This is a restraint from the tragedy of the commons and in fact a resolution to the “paradox” (Zea-Cabrera et al. 2006*b*,*a*).

Zea-Cabrera et al. (2006*b*) also found that spatial structure restrains competition with a model that fully separates individuals but implements a mixing of the water availability across sites. Here, although I unrealistically assume the water and nitrogen does not move across sites, the overlapping individual root zones effectively serve to mix the resource availability. These different implementations of “mixing” in these two models achieve different things, however. In the model presented here, mixing is not an extrinsic environmental parameter but rather an emergent property of the evolutionary optimization within the plant community. So here we find the level of resource sharing determined by the biology itself. In a way, this compensates for the lack of realism in assuming resources do not diffuse across sites. If that assumption is relaxed, plants will only influence one another’s resource availability from greater distances. Distances need to send roots (*x*_ESS_) will decrease, but the qualitative results of the influence of competition will hold – the greater the number of individuals interacting, the weaker the incentives are for fine-root overproliferation.

From the second, “hex-grid competition” model we find that the most successful plants are those that do not send their roots as far as possible, but keep them close to home while still sharing space with neighbors. Competing with as few individuals as possible (while still competing) is the best strategy, independent of the distance to neighbors or the cost of moving roots through soil (if leakage Δ is zero; if Δ is positive, see exceptions below). Understanding this result of the second model is aided by the predictions of the first. From the first model, we find that incentives for competitive overinvestments are strongest when individuals are competing with the fewest individuals. Communities suffer, but individuals receive the largest payoffs to overinvestment in fine roots when there are fewer competitors (figure 2). This is because, if an individual invests more in roots than its neighbors, it gets two benefits: first, an increase in water and/or nitrogen uptake from the additional fine roots, and second, an increase in relative water uptake, or relative fitness, due to the effect of the roots on decreasing neighbors’ water uptake. The second benefit is greater with fewer competitors. The negative impact of competitive roots on each neighbor is higher, increasing the mutant’s relative fitness further. In the full spatial grid of the second model, individuals that constrain their roots to intermingling with only the closest neighbors’ roots have higher relative fitness and dominate over evolutionary time.

Other instances of restrained belowground competition in competitive optimization models have been found. O’Brien et al. (2007) investigated optimal root placement for two individuals in competition. Using a more finely discretized treatment of space, they find that the individuals’ root systems will overlap, but not entirely. This model differed from ours however, as individuals were also allowed to explore competitor free space. In our treatment here, every spot is the home space of some individual, we find this restraint in a preference for home soil 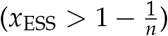 and extension to only those closest neighbors.

With the second model, we also find cases in which the best strategy is not to compete with only the closest neighbors. Specifically, when there are no costs of competition, the best strategy for plants is to spread roots evenly over all available space (just as in the *n*-competitor model). To the opposite extreme, there are cases in which the best strategy is to keep all roots at home. This occurs if leakage loss of the limiting resource drives high fine-root investment for an individual foraging, ignoring the threat of competition from neighbors. This happens when the first-order leakage loss rate is high (Δ), and resources are low (*R* or *N*, depending on *q*) or the cost of moving roots through space is high (*c*_x,0_, figures 4). Figure C5 shows these instances occur when the foraging strategy and competitive optimization of fine-root allocation converge. Finally, however, if there are no resource-dependent leakage losses (Δ = 0), it is never the best strategy to keep all roots at home (figures C2 and C3).

It is interesting to note that unlike the restraint from competition caused by a cost of extending roots through space (*c*_x_), the restraint from competition caused by an increase in optimal allocation of roots for foraging (high leakage, Δ) comes without benefits to the rest of the plant (figure 4, C5). This mechanism of restrained competition is not a restraint from the tragedy of the commons, but rather, a harsh environment overall.

It is still up for debate whether leakage losses (Δ > 0) are common. It is often assumed that leakage losses of nitrogen or water from systems are evidence that those resources are not limiting (Aber et al. 1989; Porporato et al. 2002). If that is true, and leakage losses do not occur when plants are limited by belowground resources, then our model predicts plants should always be in or be growing toward a condition where their roots are intermingled with their closest neighbors.

Although direct tests of many of the mechanisms unveiled here may be difficult due to the evolutionary (long time scale) nature of the driver, there is support for the effects of these insights from various empirical studies. At the most basic level, the insight that plants should almost always be in or working toward competition with one another is supported by analyses of fine root distributions in soil, which show that the roots of several species are commonly intermingled in forests (Göttlicher et al. 2008; Casper et al. 2003) and a grassland (Frank et al. 2015). And meta-analyses show that competition can be strong even in resource poor and sparse environments (Schoener 1983; Fowler 1986; Goldberg et al. 1999). Demonstrating that belowground competition is commonly a major influence on plant community assembly, we see major changes in ecosystems when that competition is not possible. In a study of Central European forests, Hölscher et al. (2002) find that only in the rare portions of forest that exist on rock screes the dominant species, *Fagus sylvatica* gives way to a mixture of other species. Perhaps in the extremely rocky environment, developing interactions between individuals is physically more difficult. Indeed in these areas the intermingling of fine roots is much lower than the rest of the forest. With new methods for identifying fine roots to the species and possibly individual level, more direct testing of model predictions will soon be feasible (e.g Valverde-Barrantes et al. 2015).

In order to focus the investigation here, we have ignored many of the fascinating aspects of belowground plant ecology – interactions with microbial communities (Bever et al. 2009), depth distributions (Schenk and Jackson 2002), nutrient hot spots (Hodge 2004; Chen et al. 2018), hydraulic lift (Yu and D’Odorico 2015), facilitation among individuals (Tang et al. 2018; De Parseval et al. 2017), and even size variation (Goldberg et al. 2017). But, the simplified framework has allowed us to clearly understand surprising and potentially fundamental feedbacks. Initial analyses of the inclusion of these listed phenomena have not changed these basic qualitative insights. But I look forward to investigating their interactions in more detail. Such work is likely to produce intriguing and useful refinements to this understanding.

## Conclusion

Through building and analyzing strategically simple models, we develop new hypotheses of the importance and nature of belowground competition across environmental resource gradients. We find that the physical fact of a cost of growing roots horizontally through space can restrain over-proliferation of fine roots with a net benefit on aboveground biomass. Only if plants are in environments where foraging for resources belowground is so tough that competition does not matter, or they face physical barriers to interactions with one another, are there incentives for individuals to disengage from competition belowground entirely.

In almost all cases, we find individuals have incentives to compete with their closest and only their closest neighbors. This leads to the highest payoffs of overproliferation of fine roots and thus maximizes the effects of competition while minimizing the number of competitive interactions among individuals.

These insights help explain the ubiquity of the importance of competition across environmental gradients and give specific predictions for the optimal level and extent of belowground fine root extension. They also show the importance of considerations of the cost of growing roots through space for quantitative predictions of allocation strategies and thus ecosystem carbon storage (Foley et al. (1996)).

## Acknowledgments

I thank Ray Dybzinski, Norma Fowler, and Steve Pacala for helpful conversations. I thank Martin Weiser for comments that improved the clarity of the manuscript. I thank the National Institute for Mathematical and Biological Synthesis, an Institute sponsored by the National Science Foundation through NSF Award #DBI-1300426, with additional support from The University of Tennessee, Knoxville and the University of Texas at Austin for funding.

